# Influences of rare protein-coding genetic variants on the human plasma proteome in 50,829 UK Biobank participants

**DOI:** 10.1101/2022.10.09.511476

**Authors:** Ryan S. Dhindsa, Oliver S. Burren, Benjamin B. Sun, Bram P. Prins, Dorota Matelska, Eleanor Wheeler, Jonathan Mitchell, Erin Oerton, Ventzislava A. Hristova, Katherine R. Smith, Keren Carss, Sebastian Wasilewski, Andrew R. Harper, Dirk S. Paul, Margarete A. Fabre, Heiko Runz, Coralie Viollet, Benjamin Challis, Adam Platt, AstraZeneca Genomics Initiative, Dimitrios Vitsios, Euan A. Ashley, Christopher D. Whelan, Menelas N. Pangalos, Quanli Wang, Slavé Petrovski

## Abstract

Combining human genomics with proteomics is becoming a powerful tool for drug discovery. Associations between genetic variants and protein levels can uncover disease mechanisms, clinical biomarkers, and candidate drug targets. To date, most population-level proteogenomic studies have focused on common alleles through genome-wide association studies (GWAS). Here, we studied the contribution of rare protein-coding variants to 1,472 plasma proteins abundances measured via the Olink Explore 1536 assay in 50,829 UK Biobank human exomes. Through a variant-level exome-wide association study (ExWAS), we identified 3,674 rare and significant protein quantitative trait loci (pQTLs), of which 76% were undetected in a prior GWAS performed on the same cohort, and we found that rare pQTLs are less likely to be random in their variant effect annotation. In gene-based collapsing analyses, we identified an additional 166 significant gene-protein pQTL signals that were undetected through single-variant analyses. Of the total 456 protein-truncating variant (PTV)-driven *cis*-pQTLs in the gene-based collapsing analysis, 99.3% were associated with decreased protein levels. We demonstrate how this resource can identify allelic series and propose biomarkers for several candidate therapeutic targets, including *GRN, HSD17B13, NLRC4*, and others. Finally, we introduce a new collapsing analysis framework that combines PTVs with missense *cis*-pQTLs that are associated with decreased protein abundance to bolster genetic discovery statistical power. Our results collectively highlight a considerable role for rare variation in plasma protein abundance and demonstrate the utility of plasma proteomics in gene discovery and unravelling mechanisms of action.

## Introduction

Proteins are a cell’s functional unit, and changes in protein abundance can profoundly affect biological processes and human health. Genetic variation, either within or near the protein-encoding gene (*cis*) or anywhere else in the genome (*trans*), can dramatically impact protein expression, folding, secretion, and function. Moreover, most medicines exert their effects by modulating protein levels or function. Identifying genetic variants that affect protein levels (i.e., protein quantitative trait loci, or pQTLs) has the potential to elucidate disease mechanisms, reveal new drug targets, and enhance biomarker discovery.

Proteins circulating in the blood can originate from multiple organs and cell types and include actively secreted proteins and those that leak from damaged cells elsewhere in the body. The plasma proteome can thus provide a snapshot of the current state of human health.^1^ Recent advances in high-throughput aptamer- and antibody-based proteomic platforms have enabled population-scale measurements of plasma proteins. Studies integrating plasma protein measurements with genotype array data have identified thousands of associations between genetic variants and plasma protein concentrations.^2-4^ These transformational pQTL atlases have helped prioritize candidate causal genes at genome-wide association study (GWAS) loci and have revealed potential drug repositioning opportunities. However, because these studies used genotype array data, the identified pQTLs were mainly common, non-coding variants, and often confounded by correlated non-causal signals. Compared to common variants, rarer protein-coding variants tend to confer much larger biological effect sizes, but their role in influencing human plasma protein abundances remains largely unknown.

Here, we systematically evaluated the role of rare variation in plasma protein abundance by analyzing exome sequence data and plasma levels of 1,472 plasma protein abundances measured in 50,829 UK Biobank participants. We first performed variant- and gene-level association tests to identify the *cis*- and *trans*-influences of protein-coding variation on plasma protein levels across the allele frequency spectrum. We then demonstrated how the inclusion of *cis*-acting missense variants in a traditional gene-level collapsing analyses framework augments drug target discovery and validation studies.

## Results

### UKB-PPP cohort characteristics

We performed proteomic profiling on blood plasma samples collected from 54,273 UKB participants using the Olink Explore 1536 platform, which measures 1,472 protein analytes and 1,463 unique proteins. As previously described, the UKBiobank Pharma Plasma Proteome cohort (UKB-PPP) includes plasma collections from 46,673 randomly selected participants (“randomised baseline”), 6,365 individuals chosen by the UKB-PPP consortium members (“consortium-selected”), and 1,268 individuals who participated in the COVID-19 repeat imaging study at multiple visits^2^. Exome sequencing data were available for 51,545 (95%) of these 54,273 participants, which we processed through our previously published cloud-based pipeline.^5^ Through rigorous sample QC, we removed samples with low sequencing quality and from closely related individuals as previously described (**Methods**). After further quality control based on the proteomics data (**Methods**), 50,829 (94%) multi-ancestry samples were available for downstream analyses. Of these, 47,345 (87%) were of European descent.

### Protein QTL signals through ExWAS

In our previous UKB-PPP paper, we used microarray data to perform pQTL mapping for 1,463 protein assays and identified 10,248 primary genetic associations.^2^ These analyses were limited to common variants and imputed rarer variants. Here, with the availability of whole-exome sequencing data, we directly tested for associations between variants with minor allele frequencies (MAF) as low as 0.005% in individuals of European ancestry without relying on imputation. We first performed an exome-wide, variant-level pQTL association test (ExWAS) between 1,472 plasma protein abundances and 626,929 exome sequencing variants identified in 47,345 UK Biobank participants (**Fig. 1A** and **Supplementary Table 1;** Methods). We performed an n-of-one permutation analysis (2.8 billion statistical tests) to define a variant-level significance threshold as previously described.^5^ Based on this null distribution, we identified p≤1×10^−8^ as an appropriate p-value threshold (Methods, **Supplementary Table 2**). Genomic inflation was well-controlled with a median λ_GC_ of 1.04 (95% range 1.00 – 1.10) (**Supplementary Fig. 1, Supplementary Table 3**).

**Figure 1.**
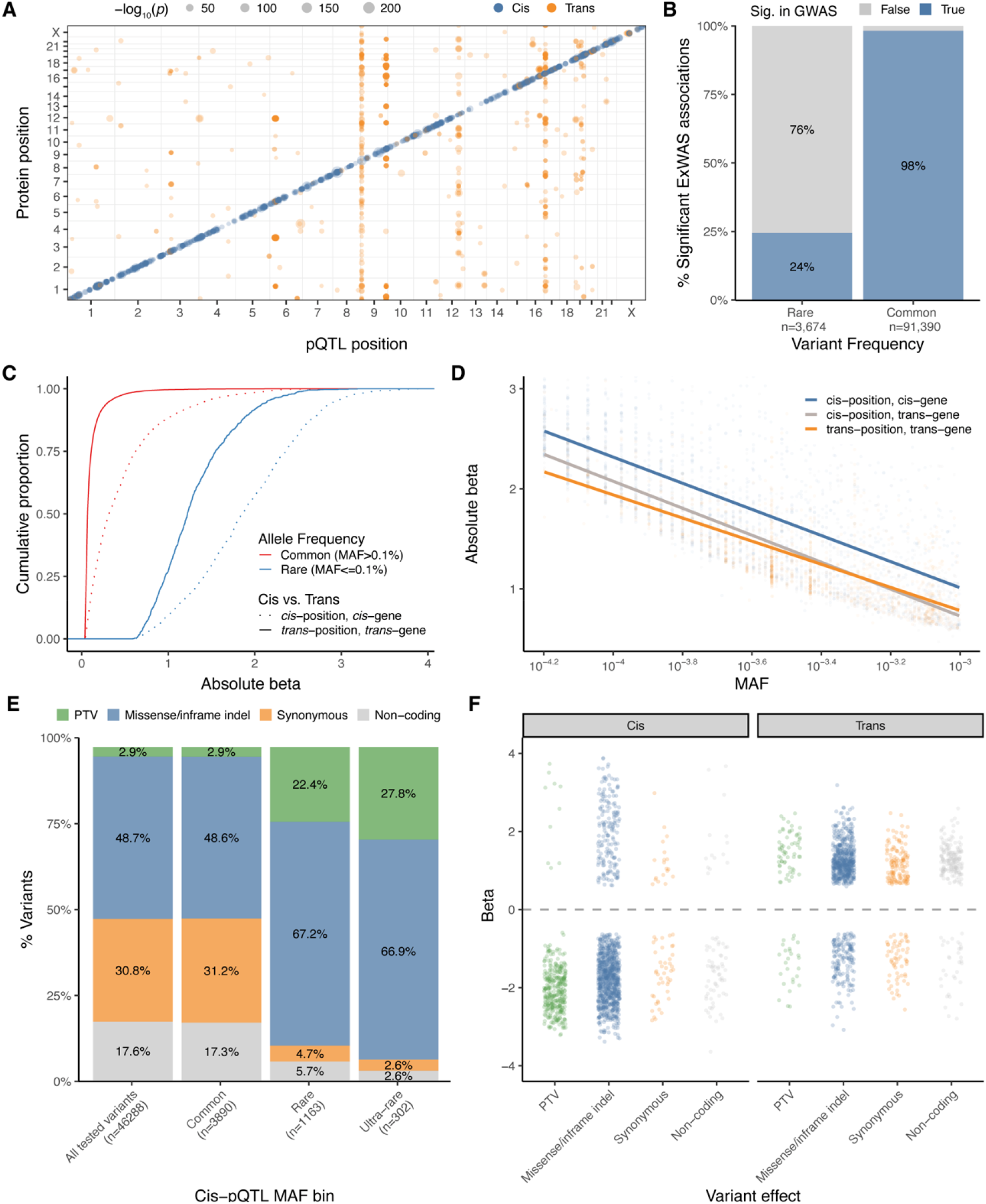
Exome-wide association study. **(A)** Summary of significant (p≤1×10^−8^) *cis* and *trans* pQTLs across the genome, limited to variants with a minor allele frequency (MAF) < 0.1%. **(B)** Percentage of significant rare (MAF≤0.1%) and common (MAF>0.1%) ExWAS pQTLs that were also significant in the UKB-PPP GWAS. **(C)** Effect size distributions of *cis-* versus *trans*-pQTLs stratified by allele frequency. **(D)** Effect sizes of rare (MAF≤0.1%) pQTLs. **(E)** The proportion of significant *cis*-pQTLs per variant class across three minor allele frequency (MAF) bins. “All tested variants” refers to the total number of variants occurring in the genes corresponding to the proteins measured via the Olink platform that were included in the ExWAS. For all plots, if the same genotype-protein association was detected in multiple ExWAS models, we retained the association with the smallest p-value. **(F)** Effect sizes of significant rare pQTLs in each variant class (PTV = protein-truncating variant).

We next compared the concordance between variant-level associations for variants included in our ExWAS that were also included in our prior GWAS,^2^ including imputed variants. The effect sizes (β) of nominally significant ExWAS protein-coding pQTLs (p<1×10^−4^) strongly correlated with the microarray-derived pQTLs (*r*^*2*^=0.96, **Supplementary Fig. 2**). Furthermore, 98% of the study-wide significant autosomal common pQTLs (MAF > 0.1%) in our study were also significant in the prior UKB-PPP GWAS (**Fig. 1B**). However, among the rare (MAF≤0.1%) autosomal pQTLs from our ExWAS analysis, only 24% were significant in the GWAS. These results illustrate the importance of exome sequencing in detecting associations for well-powered rarer variants.

We found a total of 5,355 (16.2%) coding variants that significantly affected the abundance of the encoded protein (i.e., *cis*-pQTLs). We also identified 10,768 (32.6%) coding variants that affected the abundance of any other protein that was greater than 1 megabase pair (Mbp) away from the protein directly encoded by the gene harboring the variant (i.e., *trans*-pQTLs) (**Supplementary Table 1 - ExWAS plt1×10-6**). Finally, we identified 16,887 (51.2%) *trans* pQTLs that fell within 1 Mbp of the gene encoding the protein whose level was altered, which we refer to as “*trans*-gene, *cis*-position” pQTLs. We reasoned that many *trans-*gene, *cis*-position pQTLs were contaminated by linkage disequilibrium (LD). In support of this, the relative proportion of *cis-* and *trans*-pQTLs differed among rare variants (MAF≤ 0.1%), in which 1,465 (47.3%) were *cis*-pQTLs, 592 (19.1%) were trans pQTLs, and 1,042 (33.6%) were *trans*-gene, *cis*-position pQTLs.

As purifying selection keeps variants that negatively impact fitness at low frequencies in the population, there is generally an inverse relationship between effect sizes and allele frequencies for variants that influence fitness-related traits. The median absolute effect size (β) of rare *cis-*pQTLs was 1.86, whereas the median absolute effect size of common *cis-*pQTLs was 0.32 (Wilcoxon P<10^−300^). Similarly, the absolute effect sizes of rare *trans*-pQTLs (median |β|=1.22) were significantly larger than the effect sizes of common *trans*-pQTLs (median |β|=0.07; Wilcoxon P<10^−300^) (**Fig. 1C**). Finally, even among rare variants, the effect sizes of *cis-* pQTLs (median |β| = 1.86) were greater than *trans-*pQTLs (median |β| = 1.22; Wilcoxon P=6.8×10^−125^) (**Fig. 1D**).

We next explored the number of *cis-*pQTLs per variant class across the allele frequency spectrum. Among the common *cis-*pQTLs, the proportions of PTVs, missense variants, synonymous variants, and non-coding variants closely matched the proportions observed for the total variants included in the ExWAS (i.e., the expected null distribution). In comparison, PTVs and missense variants encompassed a significantly larger percentage of rare (MAF<0.1%) and ultra-rare (MAF<0.01%) *cis-*pQTLs **(Fig. 1D, Supplementary Table 4)**. These results reinforce the observation that the common protein-coding pQTLs are more confounded by linkage disequilibrium (LD), making it challenging to confidently ascribe causality to these variants without additional experimental data.

This catalogue of protein-coding pQTLs allows us to compare the effects of different classes of protein-coding variants on protein abundances. Of the 1,465 significant rare *cis*-pQTLs, 345 (23.5%) were protein-truncating variants (PTVs), 983 (67.1%) were missense or inframe indel variants, 63 (4.3%) were synonymous variants, and 74 (5.1%) were noncoding variants (**Fig 1E; Supplementary Table 4**). As expected, nearly all the rare *cis*-pQTLs corresponding to PTVs were associated with decreased protein abundances (n=335 of 345; 97%). Of the remaining 10 *cis*-pQTL PTVs associated with increased protein abundances, five (50%) occurred in the last exon of the encoding gene, suggesting these variants may result in truncated transcripts that escape nonsense-mediated decay (NMD). Two of the 10 variants were annotated as loss of splice donor sites. Rare *cis*-pQTL missense variants and inframe indels had more variable effects, though most still decreased protein abundances (n=810/983; 82%). In comparison, among the significant rare *trans-*pQTLs, only 30% (26/87) of PTVs and 23% (159/702) of missense variants/indels were associated with decreased protein abundances.

There has been tremendous interest in identifying allelic series, in which multiple variants in a gene influence a phenotype with a range of effect sizes, to prioritise candidate drug targets.^6,7^ Missense variants are particularly valuable in discovering allelic series because they can have variable biological effects, ranging from complete or partial loss-of-function, to neutral, to gain-of-function. We thus explored how often missense variants within the same gene had similar effects on protein abundance, focusing on 117 genes with at least five rare (MAF ≤ 0.1%) missense *cis-*pQTLs. Most often, rare missense variants within the same gene had a similar effect on protein abundance. For 100 out of these 117 genes (85%), at least 75% of the significant missense pQTLs decreased protein abundance. In the remaining 17 genes, the percentage of protein-lowering missense variants ranged from 17% to 60% (**Supplementary Table 1**). However, we note that we cannot rule out epitope effects, in which a sequence variant affects antibody binding either through directly altering the binding site or changing protein structure. Consequently, such effects may also result in decreased protein abundance. However, if epitope effects had a systematic impact on missense cis-pQTL signals, we would expect to see a preferential enrichment of missense variants even among the common variant cis-pQTLs. Because we see that the variant effect proportions among the common variant cis-pQTL closely match the expected null distribution **(Fig 1E)**, it suggests that it is unlikely that epitope effects are a major driver of missense cis-pQTL signals. Nonetheless, this large catalogue of pQTLs will enable rapid hypothesis generation and validation for the identification of allelic series, which can be complemented by more targeted molecular studies.

### Protein QTL signals detected through gene-level collapsing analysis

Because the power to identify statistically significant variant-level associations decreases with MAF, we next performed gene-level collapsing analyses. In this approach, we aggregate rare variants that meet a pre-defined set of criteria (i.e., “qualifying variants” or “QVs”) in each gene and test for the aggregate effect on protein levels. Here, we used ten QV models introduced in our previous UKB phenome-wide association study (PheWAS), including one synonymous variant model that serves as an empirical negative control (**Supplementary Table 5)**. These models collectively capture genetic contributions across various genetic architectures (www.azphewas.com).^5^ Another benefit of this approach in the setting of pQTL discovery is that aggregating effects across a gene should mitigate against any potential epitope effects that might arise in the variant-level setting.

In total, we tested the association between 18,885 genes and 1,472 plasma protein levels in 47,345 individuals of European ancestry (**Supplementary Table 6**). To define an appropriate significance threshold for the collapsing analyses, we considered two different null distributions: one from an n-of-1 permutation analysis (n=276 million permutation-based statistical tests) and the other based on a synonymous variant collapsing model (i.e., empirical null; n=27.6M statistical tests) (Methods, **Supplementary Tables 7** and **8)**. Both approaches converged on a p-value threshold of p≤1×10^−8^, consistent with the ExWAS threshold **(Methods)**.

We identified 4,984 significant associations across the nine non-synonymous collapsing models (**Fig. 2A**). Of these, there were 1,330 unique gene-protein abundance associations, including 693 (52%) *cis* associations, 582 (44%) *trans* associations, and 55 (4%) *trans*-gene, *cis*-position signals. This relatively low percentage of *cis*-position, *trans*-gene associations compared to the ExWAS (4% vs. 51%) highlights the strength of rare variant collapsing analysis in mitigating contamination due to LD.

**Figure 2.**
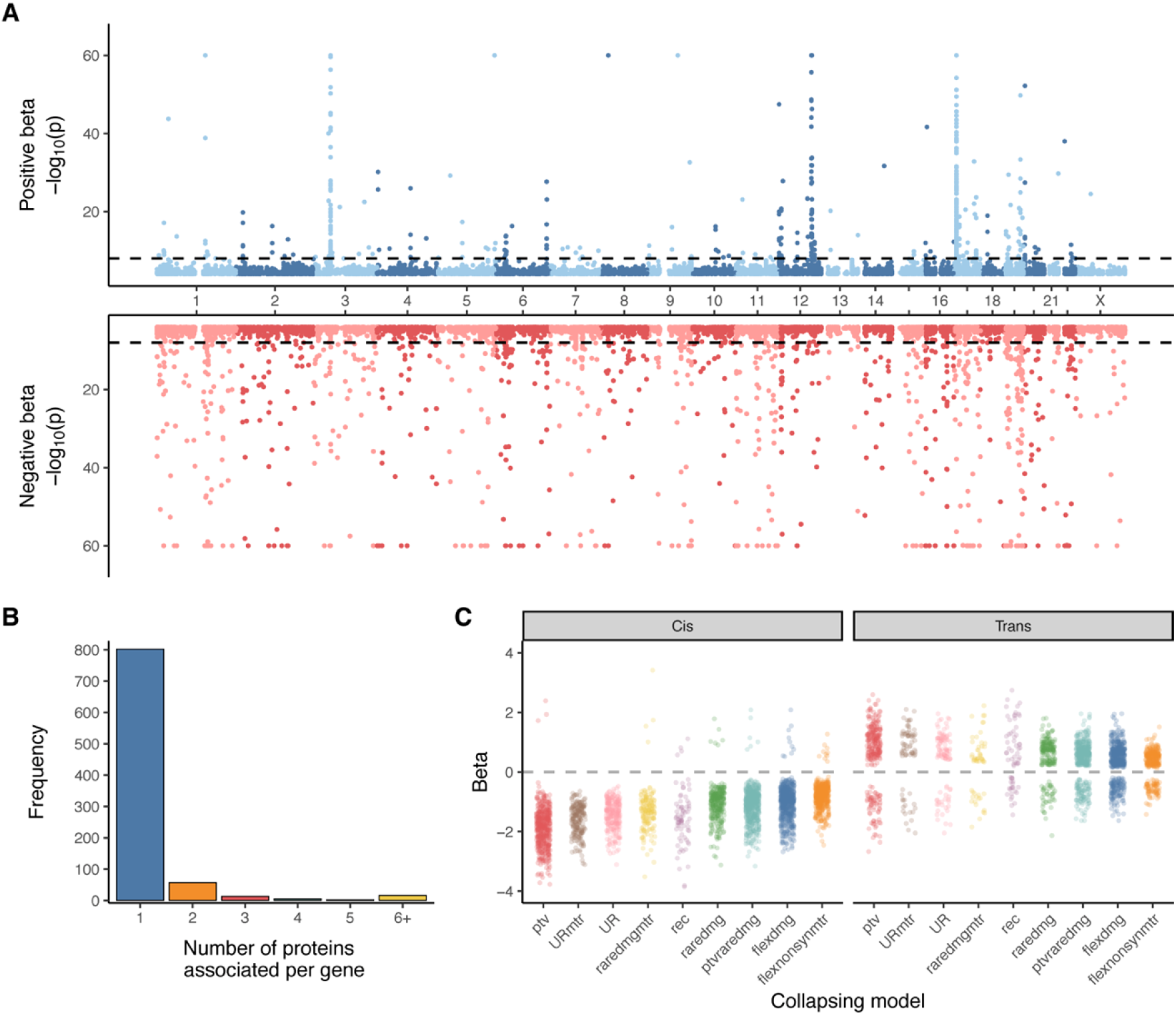
Gene-level collapsing analysis. **(A)** Miami plot of gene-protein abundance associations across nine collapsing models. We excluded the empirical null synonymous model. The y-axis is capped at 60. **(B)** The number of unique significant (p≤1×10^−8^) protein abundance associations per gene across the collapsing models. **(C)** The effect sizes of significant gene-protein associations in each collapsing model are stratified by *cis* versus *trans* effects.

Notably, 166 (12.5%) of the 1,330 gene-protein abundance signals identified via collapsing analysis did not achieve study-wide significance in the ExWAS, illustrating the increased power of this approach. Of the associations that only reached significance in the collapsing analysis, 40 (24.1%) were *cis*-pQTLs. (**Supplementary Table 6**). The greatest contribution to the 2,948 *cis*-pQTL collapsing signals came from the flexdmg model (560/2948 [19%]), followed by the ptvraredmg model (524/2948 [18%]) and the ptv model (456/2948 [15%]). In contrast to recent claims that synonymous variants are nearly as deleterious as nonsynonymous variants, we found only two significant gene-level *cis*-pQTL under the synonymous (syn) collapsing model **(Supplementary Table 8)**.^8,9^

Most pQTLs identified in the collapsing analysis were only associated with changes in abundance of a single protein (**Fig. 2B**). Among the *trans* loci, 90% of genes were associated with three or fewer proteins. However, certain genes appeared to be *trans*-pQTL “hotspots,” associated with over 20 different protein abundances. This included, *ASGR1* (n=153), *GNPTAB* (n=29), *STAB1* (n=47), and *STAB2* (n=26). ASGR1, which encodes a subunit of the asialoglycoprotein receptor, also appeared to be a *trans*-pQTL hotspot in our prior GWAS and several other large pQTL studies.^2,3,10^ *GNPTAB* encodes the alpha and beta subunits of GlcNAc-1-phosphotransferase, which selectively adds GlcNAc-1-phosphate to mannose residues of lysosomal hydrolases. The resulting mannose-6-phosphate (M6P) residues signal that the lysosomal hydrolase should be transported to the lysosome.^11^ Untagged proteins instead are secreted into the blood and extracellular space.^12^ Recessive loss-of-function mutations in *GNPTAB* are associated with Mucolipidosis III, a severe, multi-system lysosomal storage disorder (LSD) resulting in the accumulation of lysosomal substrates.^13^ Of the *GNPTAB trans-*pQTLs detected in the collapsing model, 28 (97%) are lysosomal proteins,^14,15^ 12 of which have been associated with other LSDs (**Supplementary Table 9**).^16^ Moreover, all 29 of these proteins showed increased plasma levels in PTV carriers, suggestive of reduced lysosomal targeting. Notably, there are efforts to therapeutically increase GNPTAB activity to enhance the cellular uptake of other lysosomal proteins involved in other LSDs, which could improve the efficacy of enzyme replacement therapies.^17^

Of 456 significant *cis* pQTL signals in the ptv model, 453 (99%) were associated with decreased abundance of the encoded protein, as expected. In contrast, only 54 (20%) of the 267 significant *trans* pQTL signals from the ptv model were associated with decreased protein levels. Some possible explanations for these signals include the loss of upstream regulators, reduced negative feedback, or compensatory changes. For example, we found that PTVs in *EPOR*, encoding the erythropoietin receptor, were associated with increased EPO, highlighting an example of compensatory upregulation (‘flexdmg’ model; p=3.5×10^−30^; β=0.86, 95% CI: 0.72-1.01).^18^

We observed similar patterns for the remaining collapsing models (**Fig. 2C**). Two of the collapsing models (“UR” and “URmtr”) consider ultra-rare (gnomAD MAF=0%, UKB MAF≤0.005%) PTVs and missense mutations predicted to be damaging via REVEL.^19^ The only difference between these two models is that “URmtr” only includes missense variants that fall in constrained regions of a gene based on the missense tolerance ratio (“MTR”; Methods).^20^ We compared the effect sizes between these two models to test the discriminative ability of MTR. The median absolute beta of cis loci identified through the “URmtr” model was -1.53 compared to -1.37 for the “UR” model (Wilcoxon P= 5.2×10^−7^) (**Fig. 2C**). Thus, this population genetics-based approach can effectively prioritize functional missense variants and offers a valuable layer of information on top of *in silico* pathogenicity predictors.

### Pan-ancestry collapsing analysis

Including individuals of non-European ancestry in genetic studies promotes healthcare equity and can boost genetic discovery. We performed a pan-ancestry collapsing analysis on 50,829 UK Biobank participants, including the original 47,345 European ancestry samples plus 3,484 individuals from African, Asian, and other ancestries. In this combined analysis, there were 550 unique study-wide significant gene-protein abundance associations that were not significant in the European ancestry analyses, and 163 associations that were significant in the European ancestry analyses that did not reach study-wide significance in the pan-ancestry analysis **(Supplementary Table 6)**. Of the newly significant associations, 302 (55%) were *cis*, 240 (44%) were *trans*, and 8 (1%) were *cis*-position, *trans*-gene (**Supplementary Table 10**). An example of an association that only achieved significance in the pan-ancestry analysis was the *trans* association between PTVs in *HBB* and increased levels of the monocarboxylic acid transporter encoded by *SLC16A1* (β=1.85; 95% CI: [1.33-2.37]; p=2.8×10^−12^). This association likely only reached significance in the pan-ancestry analysis due to the relative enrichment of PTVs in *HBB* variants in non-European ancestries, namely individuals of South Asian ancestry, as observed in our prior UKB exome study.^5^ Another well-known *trans* association that only became significant in the pan-ancestry analysis included PTVs in *ATM*, associated with ataxia telangiectasia and several cancers, with increased levels of alpha-fetoprotein (P=9.16×10^−9^, β=0.47, 95% CI: [0.31, 0.63]).^21^ These results add to the growing examples of how increased genetic diversity can increase power for detecting genetic associations.

### Insights into biological pathways

*Trans* associations can reflect protein-protein interactions between the encoded protein at the locus and the target protein. Several *trans* associations from the collapsing analyses capture known interactions. For example, PTVs in *PSAP*, encoding prosaposin, were associated with increased plasma abundances of progranulin (*GRN*; p=6.6×10^−17^; β=2.60, 95% CI:1.99-3.21) and cathepsin B (p=1.3×10^−11^, β=2.10, 95% CI: 1.49-2.70) (**Supplementary Table 6**). There was also a near-significant association between PTVs in *PSAP* and increased cathepsin D (p=9.5×10^−8^, β=1.61, 95% CI: 1.02-2.20). *PSAP* encodes a pro-protein that is cleaved by cathepsin D in the lysosome into four separate saposins. Recessive variants in *PSAP* are associated with various lysosomal storage disorders.^22^ Likewise, haploinsufficiency of *GRN* is associated with frontotemporal lobar degeneration (FTLD),^23,24^ and complete loss is associated with a lysosomal storage disorder called neuronal ceroid lipofuscinosis.^25^ Prior work has shown that *PSAP* (prosaposin) heterodimerizes with progranulin to regulate transport to the lysosome and regulates progranulin levels.^26,27^

Our analyses also robustly identified several *trans* associations between ligand-receptor pairs. For example, there was a significant association between nonsynonymous variants in *TSHR*, encoding the thyroid stimulating hormone receptor, and increased thyroid stimulating hormone *(TSHB)* (‘flexdmg’ model; p=2.1×10^−32^; β=0.66, 95% CI: 0.55-0.76) (**Supplementary Table 6**). Likewise, we robustly identified a *trans* association between mutations in *FLT3*, encoding the fms-related tyrosine kinase 3, and increased levels of the FLT3 ligand (FLT3LG; *‘*ptvraredmg’ model; p= 6.2×10^−21^; β=0.82, 95% CI: 0.65-0.99) (**Supplementary Table 6**). Although we highlighted well-known ligand-receptor pairs here, we anticipate that this *trans*-pQTL atlas could also help identify or suggest ligands for orphan receptors (https://astrazeneca-cgr-publications.github.io/pqtl-browser).

This resource also enables the discovery of functional biological networks. For example, we observed four rare NLRC4 protein-coding variants in the ExWAS that were associated with substantial changes in plasma levels of the proinflammatory cytokine IL-18 (**Supplementary Table 1)**. These included one frameshift variant and one missense variant associated with reduced protein levels, and two putatively gain-of-function missense variants associated with higher levels (**Table 1**). Only one of these variants was detected in our previous GWAS of the same cohort.^2^ *NLRC4* encodes the NLR family CARD domain-containing protein 4 that is involved in inflammasome activation.^28^ Prior studies have shown that rare, hypermorphic missense variants in this gene cause autosomal dominant infantile enterocolitis, characterized by recurrent flares of autoinflammation with elevated IL-18 and IL-1β levels.^29^ IL-18 has also been implicated as an inflammatory mediator of several other autoimmune diseases.^30^ We did not find any significant associations between any of these four mutations and clinically relevant phenotypes in our published phenome-wide association study of 470,000 UK Biobank exomes (https://azphewas.com).^5^ These data suggest that pharmacologic inhibition of NLRC4 may be safe. They also demonstrate that some rare putative gain-of-function mutations in this gene may not be sufficient to cause an observable phenotype, highlighting the value of this resource in clinical diagnostic settings.

**Table 1.** *NLRC4* allelic series. The four trans-pQTLs in *NLRC4* associated with changes in IL-18 levels from the ExWAS. MAF = minor allele frequency.

| *NLRC4* variant | Consequence | IL-18 beta, [95% CI] | P-value | UKB European MAF |
| --- | --- | --- | --- | --- |
| chr2:32252592:CA>C | Frameshift | -1.15, [-1.42, -0.89] | $2.2 \times 10^{-17}$ | 0.05% |
| chr2:32238296:C>A | Missense (p.Gly786Val) | -0.74 [-0.83, -0.65] | $3.2 \times 10^{-60}$ | 0.5% |
| chr2:32224523:C>A | Missense (p.Asp1009Tyr) | 2.00 [1.40, 2.59] | $5.0 \times 10^{-11}$ | 0.01% |
| chr2:32250993:C>T | Missense (p.Gly291Ser) | 1.97 [1.20, 2.74] | $5.0 \times 10^{-7}$ | 0.006% |

Beyond mapping protein regulatory pathways, this rich catalogue of protein-coding pQTLs can address several components of drug development, including the identification of novel genetic targets, discovering mechanisms of actions or biomarkers for drug targets, safety profiling, and drug repositioning opportunities. For example, there have been recent efforts to inhibit HSD17B13 based on the discovery that a splice variant (*rs72613567*) in this gene may protect against chronic liver disease.^31^ Our ExWAS revealed that this splice variant also associated with altered levels of HYAL1 (P=7.4×10^−10^, β=-0.06, 95% CI: [-0.07, 0.04]), SMPD1 (P=2.2×10^−11^, β=-0.05, 95% CI: [-0.06, -0.03]), CES3 (P=1.5×10^−12^, β=0.07, 95% CI: [0.05, 0.08]), GUSB (P=7.9×10^−9^, β=0.04, 95% CI: [0.03-0.05]), and PDGFC (P=4.8×10^−9^, β=0.04, 95% CI: [0.03, 0.06]) (**Supplementary Table 1**). Further research into the individual and combined effects of these previously undescribed relationships could help elucidate how this splice variant confers the observed reduced liver disease risk.

These vignettes provide some examples of how this expansive pQTL resource can aid many different drug discovery efforts (**Fig. 3A**). We have made the ExWAS and collapsing pQTLs publicly available through a pQTL-specific interactive portal to empower the broader research community (**Fig. 3B;** https://astrazeneca-cgr-publications.github.io/pqtl-browser).

**Figure 3.**
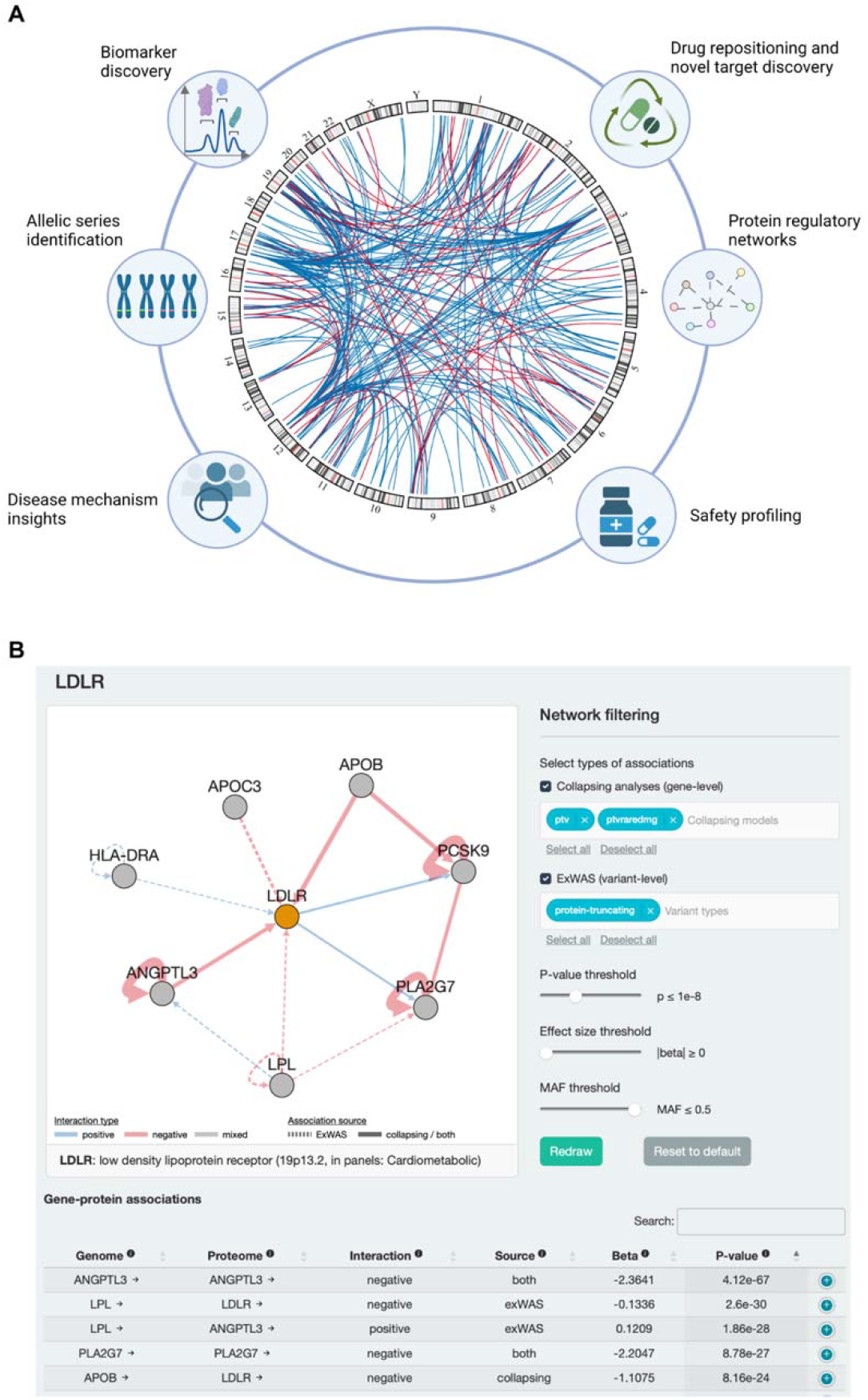
pQTL atlas and interactive browser. **(A)** Illustration of potential applications of this trans-pQTL atlas to drug development. The chord diagram represents *trans*-pQTLs detected in our collapsing analysis (p≤1×10^−8^). **(B)** The AstraZeneca pQTL browser, highlighting *LDLR* as an example user query. Users can browse pQTLs from both the ExWAS and gene-based collapsing analyses using an intuitive range of parameters and thresholds.

### Clonal haematopoiesis of indeterminate potential

The age-related acquisition of somatic mutations that lead to clonal expansion of haematopoietic stem cell populations (i.e., clonal haematopoiesis, or “CH”) has been associated with an increased risk of haematological cancer, cardiovascular disease, infection, cytopenia, and other diseases.^32,33^ To identify plasma protein changes with CH, we performed a gene-level collapsing analysis in which we defined QVs as clonal somatic variants in 15 genes recurrently mutated in myeloid cancers (see Methods) using a predefined list of variants and considered four different variant allele frequency (VAF) cut offs (S**upplementary Table 11**). In this setting, we excluded 290 individuals diagnosed with a haematological malignancy diagnosis pre-dating sample collection. We observed that the most significant (p≤1×10^−8^) associations were achieved with VAF≥ 10% cut-off (**Supplementary Table 12**). Under this model, we detected 13 *trans* protein associations with somatic mutations in *JAK2*, five with *TET2*, four with *SRSF2*, and three with *ASXL1*. Strikingly, there was no overlap between the protein abundances associated with each of these four genes, suggesting distinct downstream effects of the somatic events detected in each.

Somatic JAK2 mutations frequently cause Philadelphia-negative myeloproliferative neoplasms (including polycythaemia vera, essential thrombocythemia and primary myelofibrosis), which are associated with thromboembolic disease.^34^ Three of the *JAK2 trans-* pQTLs include proteins involved in the integrin β2 pathway, including FCGR2A, GP1BA, and ICAM2. Prior work has shown that the most common *JAK2* missense variant associated with myeloproliferative disorders (V617F) can promote venous thrombosis through activation of this pathway.^35^ The largest effect size was seen with CXCL11, encoding a chemokine.

Somatic mutations in *TET2* were associated with increased levels of the cytokine tyrosine kinase FLT3 (p=9.7×10^−15^, β=-0.50, 95% CI: [0.38, 0.63]) and decreased levels of the FLT3 ligand, FLT3LG (p=8.0×10^−54^, β=-0.95, 95% CI: [-1.01, -0.83]). FLT3 is a key regulator of hematopoietic stem cell proliferation and dendritic cell differentiation.^36^ Two other *TET2* associations included increased abundances of CD1C and CLEC4C, which are markers of conventional dendritic cells and plasmacytoid dendritic cells, respectively.^37^ Prior work has shown that roughly 30% of patients with acute myeloid leukaemia (AML) carry FLT3-activating mutations, the presence of which portend poor outcomes.^38^ There are now FLT3 inhibitors that have been found to improve survival of patients with AML.^39,40^ If the relationship between CH-*TET2* and FLT3 is causal, this could suggest potential repositioning and precision medicine opportunities.

### Augmenting PTV-driven PheWAS associations with proteomics

Understanding the functional consequences of protein-coding variants is critical to uncovering the genetic underpinnings of diseases. In the setting of gene discovery studies, it can be especially challenging to distinguish between putatively pathogenic and benign missense variants. In rare-variant aggregated collapsing analyses, researchers typically prioritise rare missense variants based on *in silico* predictions of how damaging that variant might be to the structure or function of a protein. While *in silico* scores help distinguish between neutral and potentially damaging missense variants, even the most well-performing scores only modestly correlate with experimental measures of protein function.^41^ There has thus been considerable interest in performing *in vitro* mutagenesis screens to determine the effects of many possible variants within a gene. However, the availability of protein measurements across tens of thousands of individuals can be considered a human *in vivo* mutagenesis screen since we have direct measurements of how individual observed variants impact protein levels among those carriers. We thus sought to leverage this conceptual framework in the setting of a phenome-wide association study.

In our previous rare-variant collapsing phenome-wide association study on 281,104 UKB exomes, we observed that the PTV collapsing models accounted for the greatest number of significant gene-phenotype relationships.^5^ Here, using a more extensive set of 419,387 UK Biobank exomes, we augmented our standard PTV model with missense variants associated with reduced protein abundance (i.e., ExWAS cis-pQTLs with P<0.0001; see Methods). We defined two new collapsing models: “ptvolink,” in which we included PTVs and missense pQTLs with a MAF < 0.1%, and “ptvolink2pcnt,” in which we relaxed the MAF threshold of missense variants to <2% (Methods, **Fig. 5A, Supplementary Table 5**). We tested for associations between genes encoding the Olink measured proteins and 10,017 binary and 584 quantitative phenotypes (**Supplementary Tables 13** and **14**).

**Figure 4.**
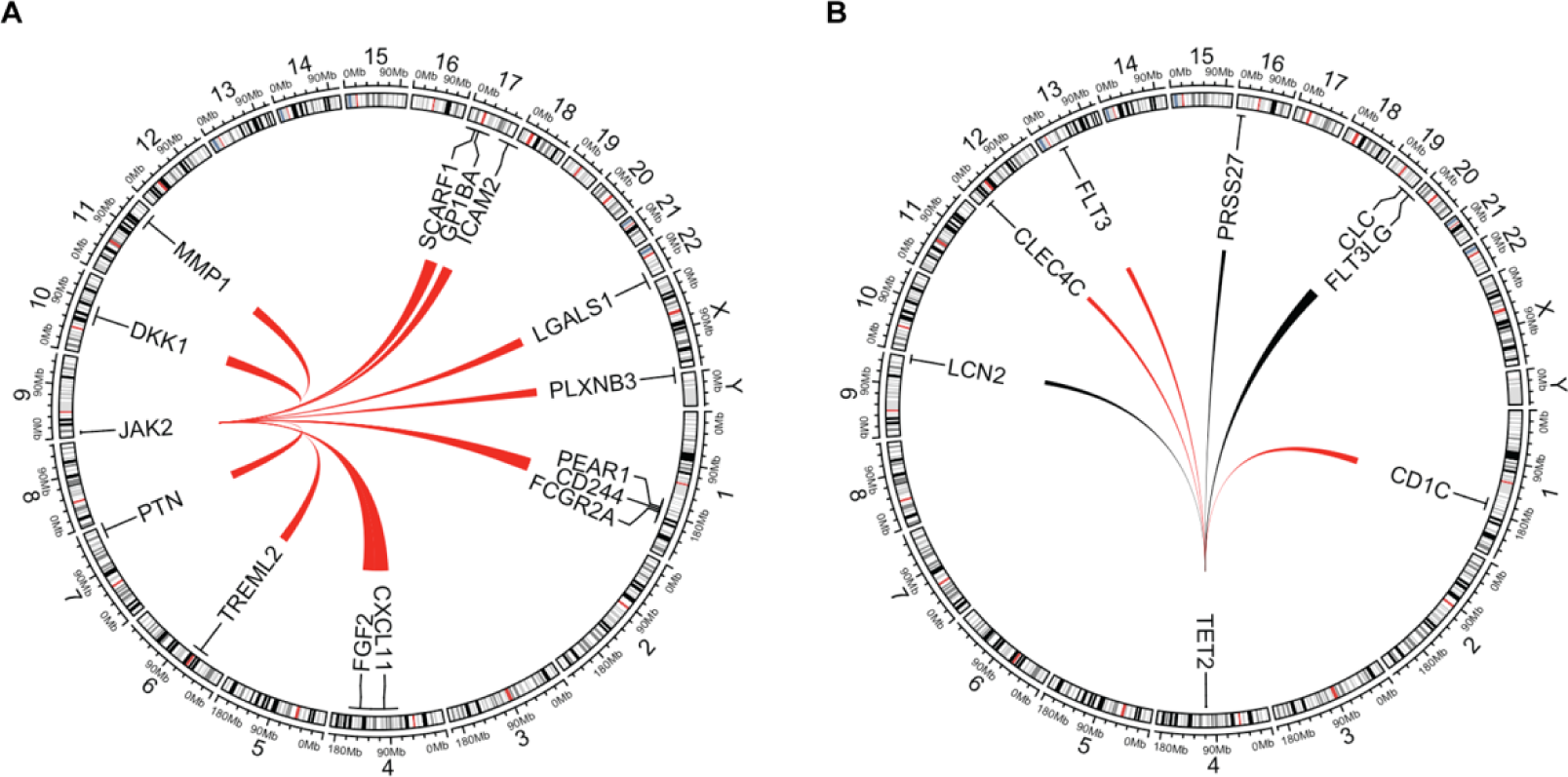
Clonal haematopoiesis *trans*-pQTL associations. **(A)** Chord diagram illustrating significant (p≤1×10^−8^) *trans*-pQTLs associated with somatic mutations in *JAK2*. **(B)** Significant *trans*-pQTLs associated with somatic mutations in *TET2*. Red lines indicate positive betas and black lines indicate negative betas. Line width is proportional to the absolute beta. For each gene, we plotted associations that were significant in any of the four collapsing models.

**Figure 5.**
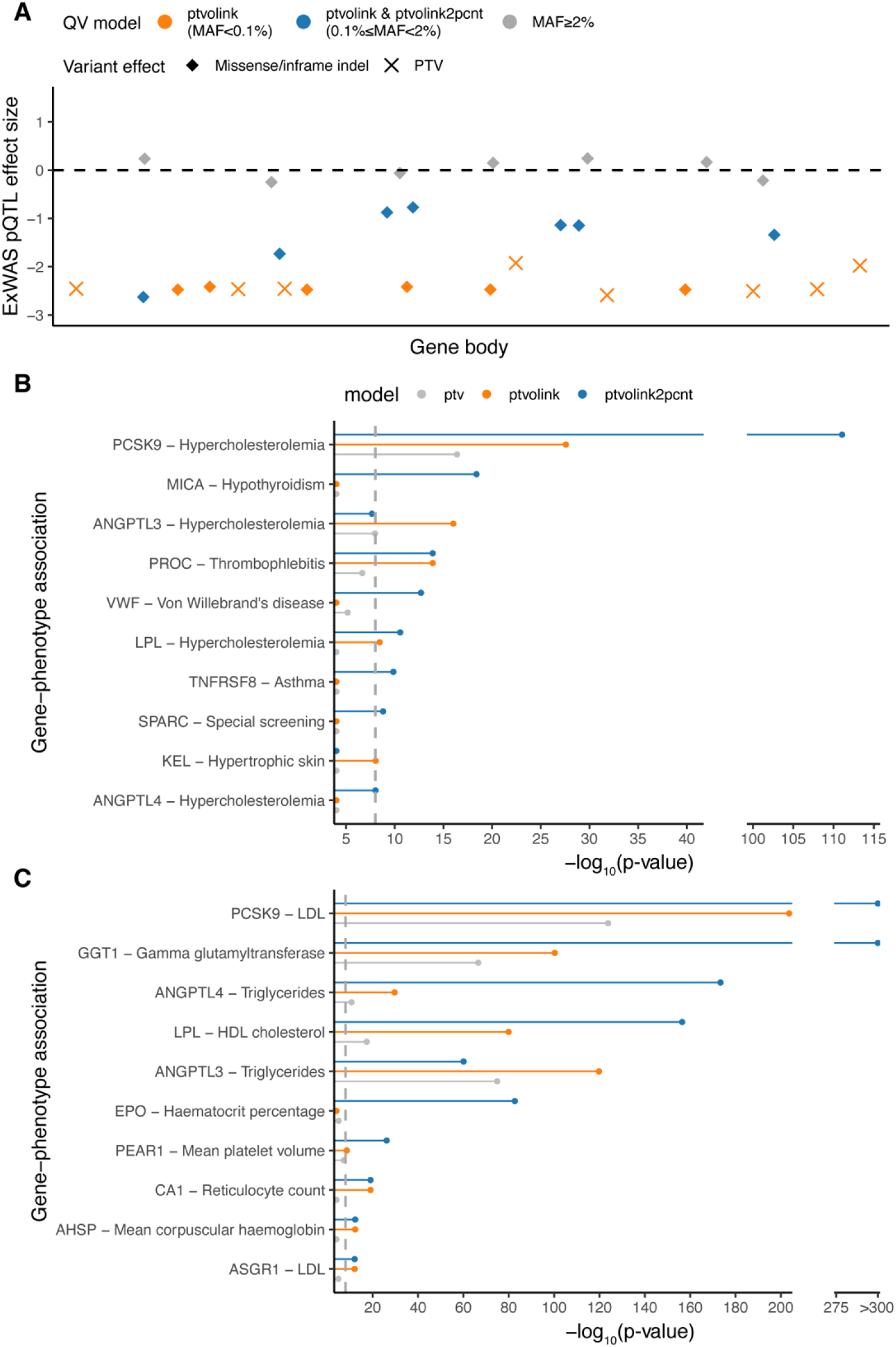
pQTL-informed collapsing analyses. **(A)** Schematic representing the pQTL-informed collapsing framework. Blue diamonds represent missense pQTLs that would be included as qualifying variants in the ptvolink model and ptvolink2pcnt model. PTVs, illustrated as X’s, are included in both models. **(B)**The p-values of gene-level associations with binary traits that improved when including PTVs and missense *cis*-pQTLs (p_ExWAS_<0.0001, ptvlolink (orange) – MAF<0.1%, ptvolink2pvnt (blue) MAF<2%). **(C)** Same as (A) but for quantitative trait associations. The x-axis is capped at 10^−300^. LDL = low density lipoprotein; HDL = high density lipoprotein. The dashed line indicates the study-wide significance threshold of p≤1×10^−8^.

The standard ptv collapsing model detected significant associations for five genes that encoded proteins measured on the Olink platform, including *ACVRL1* and *ENG* with hereditary haemorrhagic telangiectasia, *GRN* with dementia, *NOTCH1* with chronic lymphocytic leukemia, and *PCSK9* with hypercholesterolemia (P=4.0×10^−17^, OR=0.35; 95% CI: 0.27-0.46) (**Fig. 5B, Supplementary Table 15**). The significance of the well-known association between the loss of *PCSK9* and protection from hypercholesterolemia markedly improved in the pQTL-informed model (ptvolink2pcnt: P=8.7×10^−112^, OR= 0.63, 95% CI: [0.60, 0.65]). Including these missense variants, which tended to have more modest effects on protein abundance than PTVs, resulted in a weaker effect size but also clearly increased statistical power. Meanwhile, the signal of the four other gene-phenotype associations was diluted in the pQTL-informed missense models (**Supplementary Table 15)**.

Impressively, nine genes that did not achieve genome-wide significance in the standard ptv collapsing model achieved significance in at least one of the pQTL-informed models (**Fig. 5B**). The p-value of the association between *ANGPTL3* and dyslipidemia improved from 1.1×10^−8^ (OR= 0.58, 95%CI: 0.48-0.71) to 9.6×10^−17^ (“ptvolink”; OR=0.57, 95% CI: 0.50-0.66). The association between *VWF* and Von Willebrand’s disease also improved from 6.9×10^−6^ to 2.0×10^−13^. Other examples included *KEL* with hypertrophic skin disorders; *PROC* with thrombophlebitis; *LPL* with hypercholesterolemia; *MICA* with hypothyroidism, *ANGPTL4* with hypercholesterolemia; *TNFRSF8* and protection from asthma, and *SPARC* with special screening examinations (**Fig. 5B** and **Supplementary Table 15**). The second strongest association for *SPARC* was with basal cell carcinoma, suggesting that this signal arose from screening for skin cancer (ptvolink2pcnt P=4.5×10^−6^, β=2.9, 95% CI: [2.0, 4.4]).

We also identified several quantitative trait associations that increased in significance using these new collapsing models (**Fig. 5C** and **Supplementary Table 16**). Consistent with the improved p-values for related binary phenotypes, the associations between *PCSK9, ANGPTL4, LPL*, and *ANGPTL3* with lipid-related traits all improved under the ptvolink and ptvolink2pcnt models. We also found that the association between *EPO* and increased haematocrit only achieved significance in the ptvolink2pcnt model (P=2.2×10^−83^, β=-0.24, 95% CI: [-0.27, -0.22). PTVs in this gene are a well-established cause of erythrocytosis.^42^ We also detected newly significant associations between *PEAR1* (endothelial aggregation receptor) and decreased mean platelet volume (ptvolink2pcnt P=6.8×10^−27^, β=-0.26, 95% CI: [-0.31, -0.21]) and between *CA1* (carbonic anhydrase) and increased reticulocyte count (P=9.5×10^−20^, β=0.40, 95% CI: [0.32, 0.49]). Collectively, these results illustrate how including *cis* pQTLs missense variants detected through an orthogonal proteomics approach can enhance conventional loss-of-function gene collapsing analyses.

## Discussion

We performed the most extensive rare variant proteogenomics studies to date, including 1,472 plasma protein abundances measured in 50,829 UK Biobank human exomes. Our results highlight the importance of exome sequencing for rare variant associations, as most rare variant pQTLs (MAF < 0.1%) were not detected in prior GWAS. Rare *cis-* and *trans-* pQTLs conferred significantly larger effect sizes than common variant pQTLs. In the ExWAS and gene-level collapsing analysis, cis-*pQTLs* corresponding to PTVs nearly always were associated with decreased protein levels, highlighting the robustness of these associations as well as the Olink platform. Rare *trans*-pQTLs had weaker and more variable effect sizes with respect to directionality than rare *cis-*pQTLs.

We highlighted several examples of how this protein-coding pQTL atlas can address challenges in drug discovery and clinical pipelines, such as the description of an allelic series in *NLRC4* and previously undescribed plasma biomarkers for *HSD17B13*. Beyond our proof-of-concept examples, we anticipate that this resource will provide novel insights into protein regulatory networks, discovery of upstream *trans* regulators of target genes whose inhibition could increase target protein levels, performing target safety assessments, and identifying drug repositioning opportunities (**Fig. 3A**). Through our pQTL browser and our previously published UKB phenome-wide association study (PheWAS) browser (azphewas.com), researchers can now readily identify genetically anchored disease-protein abundance associations.

We additionally identified associations between somatic mutations in known CH genes and different protein abundances. Consistent with prior findings that the risks of different diseases are differentially associated across CH gene mutations, we found that each CH gene was associated with a distinct proteomic fingerprint. *TET2* associations were enriched for genes involved in dendritic cell biology, consistent with the literature association between *TET2*-CH and inflammation.

We also introduced a new gene discovery framework that incorporated missense variant *cis-*pQTLs with classical PTVs. We found that inclusion of these missense *cis-*pQTLs increased our power to detect gene-phenotype associations, particularly for genes expressed in tissues known to contribute to the plasma proteome, such as the liver. Although the p-values improved by many orders of magnitude, the effect sizes tended to be smaller in the pQTL-informed models compared to the PTV-only models, suggesting that the missense variants had less severe effects than PTVs in the corresponding genes. This collapsing framework was limited to the genes that encoded proteins included in the Olink assay. This framework could be extended to proteomics studies of other tissues or broader plasma proteome assessments in future studies.

## Methods

### UKB Cohort

The UKB is a prospective study of approximately 500,000 participants 40–69 years of age at recruitment. Participants were recruited in the UK between 2006 and 2010 and are continuously followed.^43^ The average age at recruitment for sequenced individuals was 56.5 years and 54% of the sequenced cohort comprises those of the female sex. Participant data include health records that are periodically updated by the UKB, self-reported survey information, linkage to death and cancer registries, collection of urine and blood biomarkers, imaging data, accelerometer data, genetic data, and various other phenotypic endpoints.^44^ All study participants provided informed consent.

### Olink Proteogenomics Study Cohort

Olink proteomic profiling was conducted on blood plasma samples collected from 54,273 UKB participants using the Olink Explore 1536 platform. This platform measured 1,472 protein analytes, reflecting 1,463 unique proteins measured across the four Olink panels that comprise the 1536 panel (Cardiometabolic, Inflammation, Neurology, and Oncology). The data were processed in 7 batches by Olink. Details of UKB Proteomics participant selection (across the 46,673 randomized, the 6,365 consortia selected and the 1,268 individuals participating in the COVID-19 repeat imaging study) alongside the sample handling have been thoroughly documented in Supplementary Information in Sun, et al.^2^

For WES-based proteogenomic analyses, we analysed the (95%) samples with available paired-exome sequence data. Next, we required that samples pass Olink NPX quality control as described in Sun et al. resulting in a test cohort reduction to 51,359 (95%). Given the increased variability described in Sun et al., we excluded samples in the pilot batch or with only post-COVID imaging study sampling to obtain a combined cohort of 51,291 (95%) participants. We then pruned this cohort for up to second-degree genetic relatedness (no pair with a kinship coefficient exceeding 0.1769, n= 462), resulting in 50,829 (94%) participants available for the multi-ancestry analyses performed in this paper. Europeans are the most well-represented genetic ancestry in the UKB. We identified the participants with European genetic ancestry based on Peddy^45^ Pr(EUR)>0.98 (n=47,464). We then performed finer-scale ancestry pruning of these individuals, retaining those within four standard deviations from the mean across the first four principal components, resulting in a final cohort of 47,345 (87%) individuals for the proteogenomic analyses.

### Sequencing

Whole-exome sequencing data for UKB participants were generated at the Regeneron Genetics Center (RGC) as part of a pre-competitive data generation collaboration between AbbVie, Alnylam Pharmaceuticals, AstraZeneca, Biogen, Bristol-Myers Squibb, Pfizer, Regeneron, and Takeda. Genomic DNA underwent paired-end 75-bp whole-exome sequencing at Regeneron Pharmaceuticals using the IDT xGen v1 capture kit on the NovaSeq6000 platform. Conversion of sequencing data in BCL format to FASTQ format and the assignments of paired-end sequence reads to samples were based on 10-base barcodes, using bcl2fastq v2.19.0. Exome sequences from 469,809 UKB participants were made available to the Exome Sequencing consortium in May 2022. Initial quality control was performed by Regeneron and included sex discordance, contamination, unresolved duplicate sequences, and discordance with microarray genotyping data checks.^46^

### AstraZeneca Centre for Genomics Research (CGR) bioinformatics pipeline

The 469,809 UKB exome sequences were processed at AstraZeneca from their unaligned FASTQ state. A custom-built Amazon Web Services (AWS) cloud computing platform running Illumina DRAGEN Bio-IT Platform Germline Pipeline v3.0.7 was used to align the reads to the GRCh38 genome reference and perform single-nucleotide variant (SNV) and insertion and deletion (indel) calling. SNVs and indels were annotated using SnpEFF v4.3^47^ against Ensembl Build 38.92.^48^ We further annotated all variants with their genome Aggregation Database (gnomAD) MAFs (gnomAD v2.1.1 mapped to GRCh38).^49^ We also annotated missense variants with MTR and REVEL scores.^19,20^ The AstraZeneca pipeline output files including the VCFs are available through UKB Showcase (https://biobank.ndph.ox.ac.uk/showcase/label.cgi?id=172).

### ExWAS

We tested the 626,929 variants identified in at least four individuals from the 47,345 European ancestry UKB exomes that passed both exome and Olink sample quality checks. Variants were required to pass the following quality control criteria: minimum coverage 10X; percent of alternate reads in heterozygous variants ≥ 0.2; binomial test of alternate allele proportion departure from 50% in heterozygous state P > 1 × 10^−6^; genotype quality score (GQ) ≥ 20; Fisher’s strand bias score (FS) ≤ 200 (indels) ≤ 60 (SNVs); mapping quality score (MQ) ≥ 40; quality score (QUAL) ≥ 30; read position rank sum score (RPRS) ≥ -2; mapping quality rank sum score (MQRS) ≥ -8; DRAGEN variant status = PASS; the variant site is not missing (that is, less than 10X coverage) in 10% or more of sequences; the variant did not fail any of the aforementioned quality control in 5% or more of sequences; the variant site achieved tenfold coverage in 30% or more of gnomAD exomes, and if the variant was observed in gnomAD exomes, 50% or more of the time those variant calls passed the gnomAD quality control filters (gnomAD exome AC/AC_raw ≥ 50%). In our previous UK biobank exome sequencing study we also created dummy phenotypes to correspond to each of the four exome sequence delivery batches to identify and exclude from analyses genes and variants that reflected sequencing batch effects; we provided these as a cautionary list resource for other UKB exome researchers as Supplementary Tables 25–27 in Wang et al.^5^ Since then, an additional fifth batch of exomes was released, for which we identified an additional 382 cautionary variants (**Supplementary Table 17**) on top of the original 8,365 previously described. We report the filtered-out ExWAS results from all 8,747 cautionary variants in **Supplementary Table 17**.

Variant-level pQTL p-values were generated adopting a linear regression (correcting for age, sex, age*sex, age*age, age*age*sex, PC1, PC2, PC3, PC4, batch2, batch3, batch4, batch5, batch6, batch7 and a panel specific measure of time between measurement and sampling). Three distinct genetic models were studied: genotypic (AA versus AB versus BB), dominant (AA + AB versus BB), and recessive (AA versus AB + BB), where A denotes the alternative allele and B denotes the reference allele. For ExWAS analysis, we used a significance cut-off of P≤1×10^−8^. To support the use of this threshold, we performed an n-of-1 permutation on the full ExWAS pQTL analysis. 24 of 2.8 billion permuted tests had P≤1×10^−8^ (**Supplementary Table 2**). At this P≤1 × 10^−8^ threshold, the expected number of ExWAS pQTL false positives is 24 out of the 207,409 observed significant associations (0.01%).

### Collapsing analysis

As previously described, to perform collapsing analyses we aggregated variants within each gene that fit a given set of criteria, identified as qualifying variants.^5,50,51^ In total, we performed nine non-synonymous collapsing analyses, including eight dominant and one recessive model, plus a 10^th^ synonymous variant model that serves as an empirical negative control. In each model, for each gene, the proportion of cases was compared to the proportion of controls for individuals carrying one or more qualifying variants in that gene. The exception is the recessive model, where a participant must have two qualifying alleles, either in homozygous or potential compound heterozygous form. Hemizygous genotypes for the X chromosome were also qualified for the recessive model. The qualifying variant criteria for each collapsing analysis model adopted in this study are in **Supplementary Table 5**. These models vary in terms of allele frequency (from private up to a maximum of 1%), predicted consequence (for example, PTV or missense), and REVEL and MTR scores. Based on SnpEff annotations, we defined synonymous variants as those annotated as ‘synonymous_variant’. We defined PTVs as variants annotated as exon_loss_variant, frameshift_variant, start_lost, stop_gained, stop_lost, splice_acceptor_variant, splice_donor_variant, gene_fusion, bidirectional_gene_fusion, rare_amino_acid_variant, and transcript_ablation. We defined missense as: missense_variant_splice_region_variant, and missense_variant. Non-synonymous variants included: exon_loss_variant, frameshift_variant, start_lost, stop_gained, stop_lost, splice_acceptor_variant, splice_donor_variant, gene_fusion, bidirectional_gene_fusion, rare_amino_acid_variant, transcript_ablation, conservative_inframe_deletion, conservative_inframe_insertion, disruptive_inframe_insertion, disruptive_inframe_deletion, missense_variant_splice_region_variant, missense_variant, and protein_altering_variant.

Collapsing analysis P-values were generated by using linear regression, correcting for age and sex. For all models, we applied the following quality control filters: minimum coverage 10X; annotation in CCDS transcripts (release 22; approximately 34 Mb); at most 80% alternate reads in homozygous genotypes; percent of alternate reads in heterozygous variants ≥ 0.25 and ≤ 0.8; binomial test of alternate allele proportion departure from 50% in heterozygous state P > 1 × 10−6; GQ ≥ 20; FS ≤ 200 (indels) ≤ 60 (SNVs); MQ ≥ 40; QUAL ≥ 30; read position rank sum score ≥ −2; MQRS ≥ −8; DRAGEN variant status = PASS; the variant site achieved tenfold coverage in ≥ 25% of gnomAD exomes, and if the variant was observed in gnomAD exomes, the variant achieved exome z-score ≥ −2.0 and exome MQ ≥ 30.

The list of 18,885 studied genes and corresponding coverage statistics of how well each protein-coding gene is represented across all individuals by the exome sequence data is available in **Supplementary Table 19**. Moreover, we had previously created dummy phenotypes to correspond to each of the five exome sequence delivery batches to identify and exclude from analyses 46 genes that were enriched for exome sequencing batch effects; these cautionary lists were made available in Supplementary Tables 25–27 of Wang et al 2021.^5^ Gene-based pQTL p-values were generated adopting a linear regression (correcting for age, sex, age*sex, age*age, age*age*sex, PC1, PC2, PC3, PC4, batch1, batch2, batch3, batch4, batch5, batch6, and batch7). For the pan-ancestry analysis we included additional categorical covariates to capture broad ancestry (European, African, East Asian, and South Asian).

For gene-based collapsing analyses, we used a significance cut-off of P≤1×10^−8^. To support the use of this threshold, we ran the synonymous (empirical null) collapsing model and found only five events achieved a signal below this threshold. Moreover, we performed an n-of-1 permutation on the full collapsing pQTL analysis. Only 3 of 276 million permuted tests had P≤1×10^−8^ (**Supplementary Table 7**). At this P≤1×10^−8^ threshold, the expected number of collapsing pQTL false positives is 3 out of the 4,984 (0.06%) observed significant associations.

### Down-sampled analysis

To test the robustness of the ExWAS and collapsing analysis pQTLs, we compared the correlation between the p-values derived from the full cohort to a down-sampled subset of 40,567 samples and observed very strong correlations (**Supplementary Figure 3**).

### Phenotypes

We studied two main phenotypic categories: binary and quantitative traits taken from the April 2022 data release that was accessed on 6 April 2022 as part of UKB applications 26041 and 65851. To parse the UKB phenotypic data, we adopted our previously described PEACOCK package, located at https://github.com/astrazeneca-cgr-publications/PEACOK.^5^

The PEACOK R package implementation focuses on separating phenotype matrix generation from statistical association tests. It also allows statistical tests to be performed separately on different computing environments, such as on a high-performance computing cluster or an AWS Batch environment. Various downstream analyses and summarizations were performed using R v3.6.1 https://cran.r-project.org. R libraries data.table (v1.12.8; https://CRAN.R-project.org/package=data.table), MASS (7.3-51.6; https://www.stats.ox.ac.uk/pub/MASS4/), tidyr (1.1.0; https://CRAN.R-project.org/package=tidyr) and dplyr (1.0.0; https://CRAN.R-project.org/package=dplyr) were also used.

For UKB tree fields, such as the ICD-10 hospital admissions (field 41202), we studied each leaf individually and studied each subsequent higher-level grouping up to the ICD-10 root chapter as separate phenotypic entities. Furthermore, for the tree-related fields, we restricted controls to participants who did not have a positive diagnosis for any phenotype contained within the corresponding chapter to reduce potential contamination due to genetically related diagnoses. A minimum of 30 cases were required for a binary trait to be studied. In addition to studying UKB algorithmically defined outcomes, we studied union phenotypes for each ICD-10 phenotype. These union phenotypes are denoted by a ‘Union’ prefix and the applied mappings are available in Supplementary Table 1 of Wang et al. 2021.^5^

In total, we studied 10,017 binary and 584 quantitative phenotypes. As previously described, for all binary phenotypes, we matched controls by sex when the percentage of female cases was significantly different (Fisher’s exact two-sided P < 0.05) from the percentage of available female controls. This included sex-specific traits in which, by design, all controls would be the same sex as cases.^5^ All phenotypes and corresponding chapter mappings for all phenotypes are provided in **Supplementary Table 7**.

### Detecting clonal haematopoiesis somatic mutations

To detect putative clonal haematopoiesis, somatic variants we used the same GRCh38 genome reference aligned reads as for germline variant calling, and ran somatic variant calling with GATK’s Mutect2 (v.4.2.2.0).^52^ This analysis focused on the 74 genes previously curated as being recurrently mutated in myeloid cancers.^33^ To remove potential recurrent artifacts we filtered variants using a panel of normals created from 200 of the youngest UKB participants without a haematologic malignancy diagnosis. Subsequent filtering was performed with GATK’s *FilterMutectCalls*, including the filtering of read orientation artifacts using priors generated with *LearnReadOrientationModel*.

From the variant calls, clonal somatic variants were identified using a predefined list of gene-specific variant effects and specific missense variants (**Supplementary Table 20**). Only PASS variant calls with 0.03 ≤ Variant Allele Frequency (VAF) ≤ 0.4 and Allelic Depth (AD) ≥ 3 were included. For each gene we validated the identified variants collectively as somatic by inspection of the age versus population prevalence profile **(Supplementary Figure 4)** and limited further analysis to a set of 15 genes.

### Implementing the 470K missense pQTL-augmented PheWAS

In this study, we repeated our published PheWAS here adopting the now 469,809 available UK Biobank exomes and 10,017 binary endpoints alongside 584 quantitative endpoints. To determine whether novel signals could be detected after augmenting our standard ptv collapsing analysis model with *cis*-acting missense variants identified among the UKB Proteomics subset to correlate with a reduction in corresponding protein levels. We set our *cis*-pQTL missense p-value inclusion threshold to p<0.0001 from the previously described exWAS analyses and require a negative *cis*-acting beta. We identified 3,093 missense variants with *cis*-acting negative betas (p<0.0001) among the genes encoding the 1,472 Olink protein analytes. 919 (62%) distinct genes carried at least one of these 3,093 missense variants.^5^ To assess improved signal detection over the baseline ptv collapsing model, we introduced two new collapsing models “ptvolink” and “ptvolink2pcnt”. ptvolink adopts the baseline ptv collapsing model with the only deviation being the inclusion of the 3,093 missense variants that also qualify the QC and MAF criteria as adopted for the ptv collapsing model. ptvolink2pcnt is a repeat of the ptvolink collapsing model but permits missense variants with a MAF in the UK Biobank population as high as 2% as long as they were among the list of 3,093 missense variants identified to have a p<0.0001 negative beta *cis*-pQTL signals in the Olink ExWAS analyses. Full model descriptions are available in **Supplementary Table 5**. These new *cis*-pQTL missense ptv augmented collapsing models were then compared to the standard collapsing models.

There may be instances where reduced protein levels reflect a disruption of antibody binding rather than a true biological signal. In the setting of collapsing analysis, in which we aggregate many variant effects in a gene, we expect these events to represent only a modest fraction of a gene’s complete allelic series. Moreover, in the context of this assessment, the inclusion of missense pQTLs would be expected to act conservatively (i.e., diluting the value of including such missense in the PTV proteogenomic-augmented PheWAS collapsing analyses).

The UK Biobank exomes cohort that was adopted for this refreshed PheWAS analysis was sampled from the available 469,809 UK Biobank exome sequences. We excluded from analyses 118 (0.025%) sequences that achieved a VerifyBAMID freemix (contamination) level of 4% or higher,^53^ and an additional five sequences (0.001%) where less than 94.5% of the consensus coding sequence (CCDS release 22) achieved a minimum of tenfold read depth.^54^

Using exome sequence-derived genotypes for 43,889 biallelic autosomal SNVs located in coding regions as input to the kinship algorithm included in KING v2.2.3,^55^ we generated pairwise kinship coefficients for all remaining samples. We used the ukb_gen_samples_to_remove() function from the R package ukbtools v0.11.3^56^ to choose a subset of individuals within which no pair had a kinship coefficient exceeding 0.1769, to exclude predicted first-degree relatives. For each related pair, this function removes whichever member has the highest number of relatives above the provided threshold. Through this process, an additional 24,116 (5.1%) sequences were removed from downstream analyses. We predicted genetic ancestries from the exome data using peddy v0.4.2 with the ancestry labeled 1,000 Genomes Project as reference.^45^ Of the 445,570 remaining UKB sequences, 24,790 (5.3%) had a Pr(European) ancestry prediction of less than 0.95. Focusing on the remaining 420,780 UKB participants, we further restricted the European ancestry cohort to those within ±4 s.d. across the top four principal component means. This resulted in 419,387 (89.3%) participants of European ancestry who were included in these *cis*-pQTL modified analyses.

To remove potential concerns of circularity we repeated the above ptvolink and ptvolink2pcnt collapsing model PheWAS; however, this time we removed UK Biobank participants from the PheWAS analyses if they were part of the UKB Proteomics cohort of 47,345 individuals adopted to select the 3,093 *cis*-pQTL missense variants. These results are reflected in ptvolinknoppp and ptvolink2pcntnoppp outputs (**Supplementary Table 21**).

## Acknowledgments

We thank the participants and investigators of the UK Biobank study who made this work possible (Resource Application Number 26041 and 65851). We are grateful to the research & development leadership teams at the thirteen participating UKB-PPP member companies (Alnylam Pharmaceuticals, Amgen, AstraZeneca, Biogen, Bristol-Myers Squibb, Calico, Genentech, Glaxo Smith Klein, Janssen Pharmaceuticals, Novo Nordisk, Pfizer, Regeneron, and Takeda) for funding the study. We thank the Legal and Business Development teams at each company for overseeing the contracting of this complex, precompetitive collaboration, with particular thanks to Erica Olson of Amgen, Andrew Walsh of GSK, and Fiona Middleton of AstraZeneca. We thank the UKB Exome Sequencing Consortium (UKB-ESC) members: AbbVie, Alnylam Pharmaceuticals, AstraZeneca, Biogen, Bristol-Myers Squibb, Pfizer, Regeneron and Takeda for funding the generation of the data, and Regeneron Genetics Center for completing the sequencing and initial quality control of the exome sequencing data. We are also grateful to the AstraZeneca Centre for Genomics Research Analytics and Informatics team for processing and analysis of sequencing data.

## Data availability

Association statistics generated in this study are publicly available through our AstraZeneca Centre for Genomics Research (CGR) PheWAS Portal (http://azphewas.com/) and our pQTL browser (https://astrazeneca-cgr-publications.github.io/pqtl-browser). All whole-exome sequencing data described in this paper are publicly available to registered researchers through the UKB data access protocol. Exomes can be found in the UKB showcase portal: https://biobank.ndph.ox.ac.uk/showcase/label.cgi?id=170. The Olink proteomics data are also available under dataset #[dataset ID and URL on publication depending on time of official publication]. Additional information about registration for access to the data is available at http://www.ukbiobank.ac.uk/register-apply/. Data for this study were obtained under Resource Application Number 26041.

## Code availability

PheWAS and ExWAS association tests were performed using a custom framework, PEACOK (PEACOK 1.0.7). PEACOK is available on GitHub: https://github.com/astrazeneca-cgr-publications/PEACOK/.

## Ethics declarations

The protocols for the UK Biobank are overseen by The UK Biobank Ethics Advisory Committee (EAC); for more information see https://www.ukbiobank.ac.uk/ethics/ and https://www.ukbiobank.ac.uk/wp-content/uploads/2011/05/EGF20082.pdf.

