## Supplementary Information for "Influences of rare protein-coding genetic variants on the human plasma proteome in 50,829 UK Biobank participants"

[Supplementary Table 1 – ExWAS (p<1e-8)	7](#_Toc116209494)

### Supplemental Figure 1

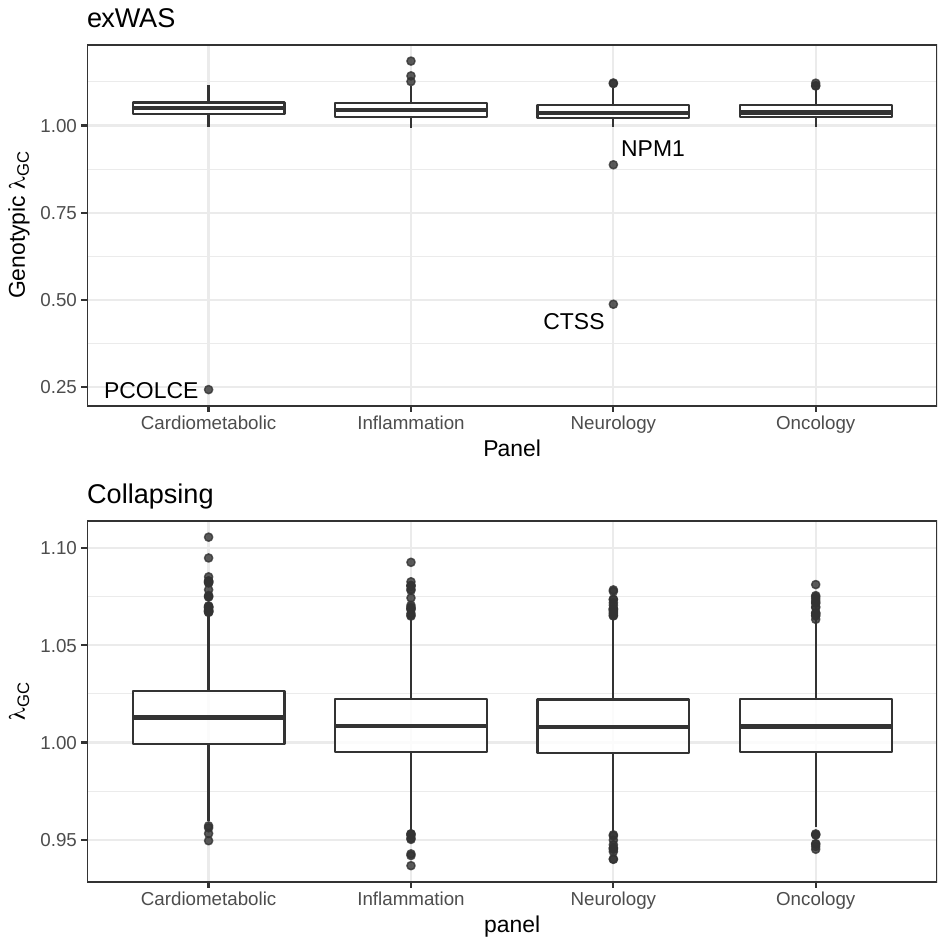

**Lambda genomic control boxplots by Olink panel.** Lambda_gc_ was computed using an n of 1 permutation approach for **(**A) exWAS genotypic model, outlying assays showing evidence of deflation (Lambda_gc_ < 1) are labelled, and relate to significant assay failures (B) collapsing analysis.

### Supplementary Figure 2

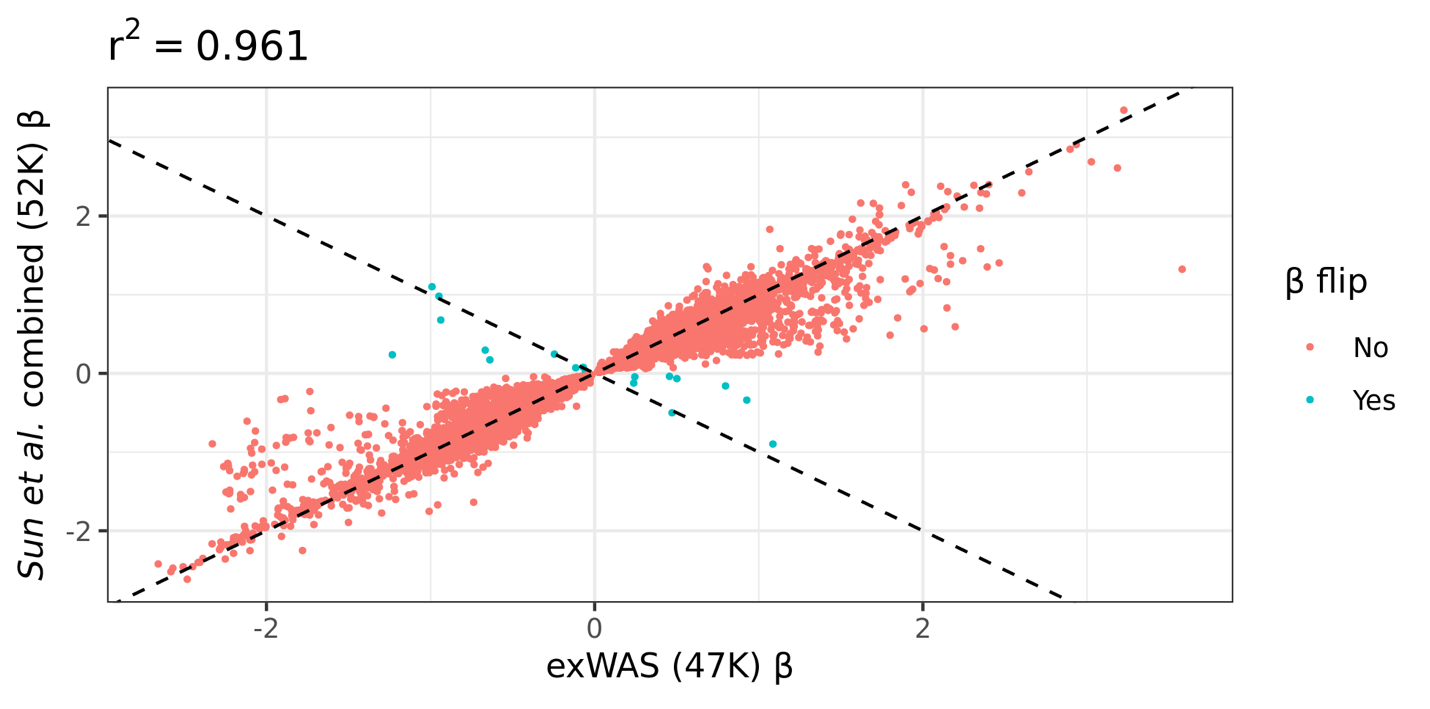

**Correlation of effect sizes between overlapping pQTLs between *Sun et al.* combined and exWAS.** Effect sizes are plotted for variants reaching p<1e-4 in both *Sun et al.* combined analysis and exWAS (genotypic model). Color indicates whether sign of the effect size is discordant between studies. Dotted lines represent lines of equivalence. The correlation coefficient reflects Pearson’s *r^2^*

### Supplementary Figure 3

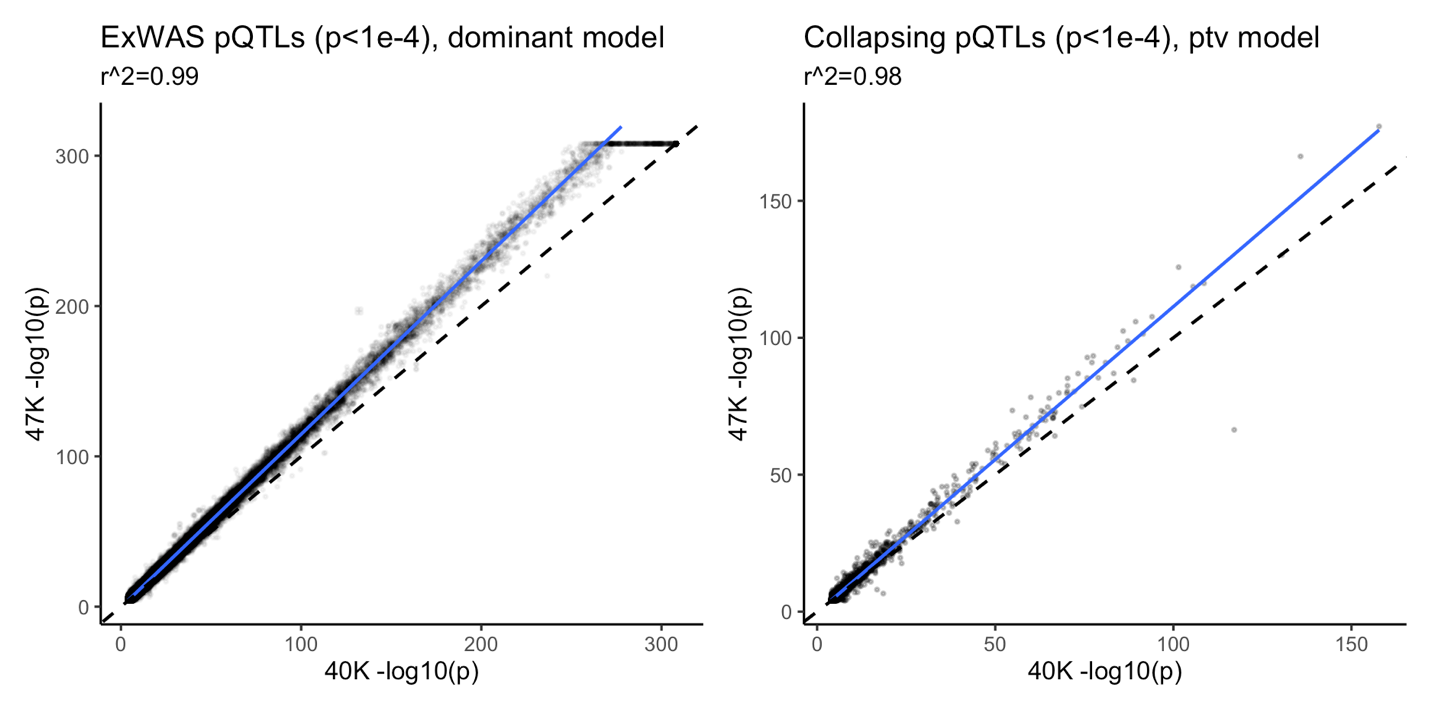

**Correlation between pQTLs detected in 40K and 47K individuals. (A)** Scatter plot of -log10(p-values) for pQTLs detected in the dominant ExWAS model for tthe full cohort (“47K”) versus the downsampled cohort (40K). We included pQTLs with a p<0.0001 (n=281,228 variants). The axes are capped at 300. **(B)** Scatter plot of p-values from the ptv model in the gene-level collapsing analysis for gene-protein associations with p<0.0001 (n=4,137). Correlation coefficients were calculated via Pearson’s R.

### Supplementary Figure 4

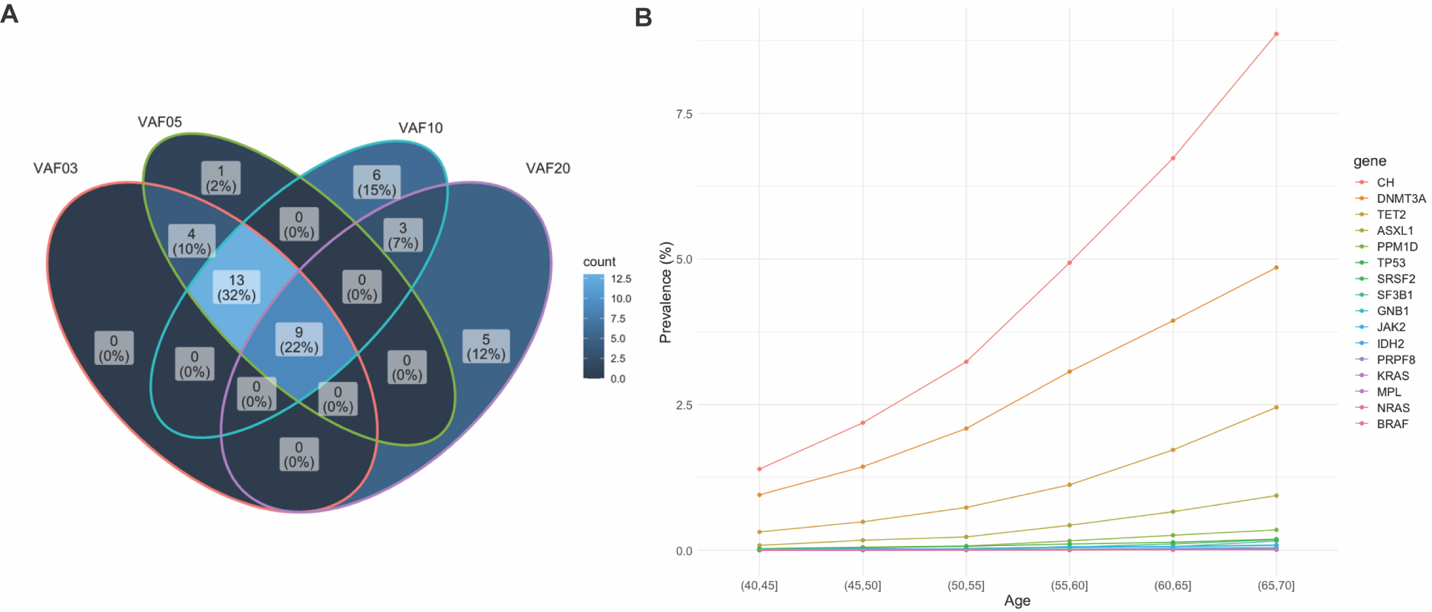

**Clonal haematopoiesis of indeterminate potential. (A)** Venn diagram illustrating the pQTL associations detected at different variant allele frequency (VAF) cut-offs (Methods). **(B)** The prevalence of putative somatic variants by age in 15 established leukemogenic genes. “CH” refers to variants observed in all 15 genes.

### Supplementary Table 1 – ExWAS (p<1e-8)

Provided as external file. List of ExWAS variants that achieved a p-value≤1x10^-8^. P-values were generated via linear regression. Cis_trans_position column refers to whether the pQTL was cis, trans, or “cis-position, trans-gene.” “Pan_p” and “Pan_beta” are the p-value and β values from the pan-ancestry analysis. All other statistics refer to the European-only analysis.

### Supplementary Table 2 – ExWAS permutation

Provided as external file. Results from the n-of-1 permutation analysis for the ExWAS (Methods).

### Supplementary Table 3 – lambda dist.

Provided as external file. Lambda distributions for the ExWAS and collapsing analyses.

### Supplementary Table 4 – ExWAS tallies

Provided as external file. Number of ExWAS variants per variant effect class. The number of study-wide significant cis-pQTLs versus the total number of tested variants binned by functional category and minor allele frequency (MAF) stratum. For each variant-protein abundance association, we retained the most significant association across the three ExWAS models (i.e., genotypic, dominant, and recessive). The P-values reflect Binomial exact tests (alternative hypothesis = greater) comparing the observed versus expected proportion of variants in each cell of the table. Variants were grouped into bins based on the following SnpEff annotations:

PTV – stop gain/loss: {stop_gained, start_lost, stop_lost}

PTV – frameshift: {frameshift_variant, bidirectional_gene_fusion, exon_loss_variant, gene_fusion}

PTV – canonical splice: {splice_donor_variant, splice_acceptor_variant}

Inframe Indels: {conservative_inframe_insertion, conservative_inframe_deletion, disruptive_inframe_deletion, disruptive_inframe_insertion}

Missense: {missense_variant, missense_variant&splice_region_variant}

Synonymous: {synonymous_variant, synonymous_variant&splice_region_variant, stop_retained_variant, initiator_codon_variant, stop_retained_variant&splice_region_variant}

Noncoding: {splice_region_variant, non_coding_transcript_exon_variant, 3_prime_UTR_variant, 5_prime_UTR_variant, 5_prime_UTR_premature_start_codon_gain_variant, intragenic_variant,

5_prime_UTR_truncation}

### Supplementary Table 5 – Collapsing models

Provided as external file.

* reflects the gnomAD global_raw MAF unless otherwise specified.
^ reflects the maximum proportion of UKB exome sequences permitted to either have ≤ 10-fold coverage at variant site or carry a low-confidence variant that did not meet one of the quality-control thresholds applied to collapsing analyses (see methods).
^#^ reflects collapsing models newly introduced compared to Wang et al. (Nature 2021).

### Supplementary Table 6 – Collapsing pQTLs

Provided as external file. List of gene-level collapsing analysis associations with a p-value ≤ 1x10^-8^. P-values were generated via linear regression. Cis_trans_position column refers to whether the pQTL was cis, trans, or “cis-position, trans-gene.” “Pan_p” and “Pan_beta” are the p-value and β values from the pan-ancestry analysis. All other statistics refer to the European-only analysis.

### Supplementary Table 7 – Collapsing permutation

Provided as external file. Results from the n-of-1 permutation analysis for the gene-level collapsing analysis (Methods).

### Supplementary Table 8 – Collapsing syn. null

Provided as external file. Results from the synonymous collapsing analysis model (p<1x10^-7^).

### Supplementary Table 9 – GNPTAB trans-pQTLs

Provided as external file. *Trans-*pQTLs for *GNPTAB* from the ptv collapsing analysis model (p≤1x10^-8^). We note whether the *trans* protein is a known lysosomal gene in the “Lysosomal?” column. The OMIM gene column indicates whether the protein is known to be associated with a lysosomal storage disease in OMIM.

### Supplementary Table 10 – multi-ancestry collapsing pQTLs

Provided as external file. List of gene-level collapsing analysis associations with a p-value < 1x10^-4^ in the pan-ancestry collapsing analysis models.

### Supplementary Table 11 – CHIP models

Provided as external file. List of qualifying variant models used in the CHIP analysis.

### Supplementary Table 12 – CHIP pQTLs

Provided as external file. List of pQTLs observed in the CHIP collapsing analysis models (p<1x10^-4^).

### Supplementary Table 13 – Studied binary phenotypes

Provided as external file. List of binary traits included in the phenome-wide association study.

### Supplementary Table 14 – Studied quant. phenotypes

Provided as external file. List of quantitative traits included in the phenome-wide association study.

### Supplementary Table 15 – Binary PheWAS

Provided as external file. Results of gene-phenotype associations from the pQTL-informed PheWAS analysis (p<1x10^-4^) for binary phenotypes.

### Supplementary Table 16 –Quant PheWAS

Provided as external file. Results of gene-phenotype associations from the pQTL-informed PheWAS analysis (p<1x10^-4^) for quantitative traits.

### Supplementary Table 17 – Cautionary variants

Provided as external file. Variant IDs for variants that we observed to have a batch effect (Methods).

### Supplementary Table 18 – Cautionary variant ExWAS

Provided as external file. ExWAS results for the cautionary variants included in Supplemental Table 17.

### Supplementary Table 19 – Gene coverage

Provided as external file. Average coverage for each protein-coding gene across all individuals in the cohort.

### Supplementary Table 20 – CHIP variants

Provided as external file. List of pre-defined CHIP variants.

### Supplementary Table 21 – noppp_ptvolink

Provided as external file. Results from the pQTL-informed PheWAS, excluding individuals who were included in the PPP cohort.

### AstraZeneca Genomics Initiative banner contributors

Rasmus Ågren, Lauren Anderson-Dring, David Baker, Carl Barrett, Maria Belvisi, Mohammad Bohlooly-Y, Lisa Buvall, Niedzica Camacho, Lisa Cazares, Sophia Cameron-Christie, Benjamin Challis, Suzanne Cohen, Regina F Danielson, Shikta Das, Andrew Davis, Sri Vishnu Vardhan Deevi, Brian Dougherty, Zammy Fairhurst-Hunter, Manik Garg, Benjamin Georgi, Carmen Guerrero Rangel, Carolina Haefliger, Mårten Hammar, Richard N. Hanna, Pernille B.L. Hansen, Jennifer Harrow, Ian Henry, Sonja Hess, Fengyuan Hu, Xiao Jiang, Kousik Kundu, Zhongwu Lai, Mark Lal, Glenda Lassi, Margarida Lopes, Kieren Lythgow, Meeta Maisuria-Armer, Ruth March, Carla Martins, Karine Megy, Rob Menzies, Erik Michaëlsson, Fiona Middleton, Daniel Muthas, Abhishek Nag, Sean O’Dell, Yoichiro Ohne, Henric Olsson, Amanda O’Neill, Kristoffer Ostridge, William Rae, Arwa Raies, Anna Reznichenko, Xavier Romero Ros, Maria Ryaboshapkina, Hitesh Sanganee, Ben Sidders, Mike Snowden, Helen Stevens, Ioanna Tachmazidou, Christina Underwood, Anna Walentinsson, Qing-Dong Wang, Andrew Whittaker, Ahmet Zehir, Jorge Zeron, Zoe Zou
